# Measuring physiological responses to stressors using a novel *Hmox1* reporter mouse

**DOI:** 10.1101/098467

**Authors:** Michael McMahon, Shaohong Ding, Lourdes P Acosta-Jimenez, Tania G. Frangova, Colin J. Henderson, C. Roland Wolf

**Author notes:** This work was funded by a European Research Council Advanced Investigator Award (number 294533) and a CRUK Programme grant (C4639/A10822), both awarded to CRW. Corresponding author: C. Roland Wolf.

## Abstract

Hmox1 protein holds great promise as a biomarker of organismal health as it is highly expressed in stressed or damaged cells. However, documenting Hmox1 expression patterns has thus far only been feasible in simple model organisms with limited relevance to humans. We now report a new Hmox1 reporter line that makes it possible to access this information in mice, the premiere model system for studying human disease and toxicology. Using a state-of-the-art strategy, we express multiple complementary reporter molecules from the murine *Hmox1* locus, including firefly luciferase to allow long-term, non-invasive imaging of Hmox1 expression, and β-galactosidase for high-resolution mapping of expression patterns *post-mortem*. We validate the model by confirming the absence of haplo-insufficiency effects, the fidelity of reporter expression, and its responsiveness to oxidative and inflammatory stimuli. In addition to providing blueprints for Hmox1 expression in mice that provide novel biological insights for follow-up, this work paves the way for the broad application of this model to regulatory toxicology studies and, also, pre-clinical investigations into human degenerative diseases.

## INTRODUCTION

Across all kingdoms of life, organisms adapt to environmental stress by inducing key evolutionarily conserved genes involved in damage prevention, mitigation, and repair (1,2). Classic examples of these cytoprotective genes are those encoding heat-shock- and DNA repair proteins, and detoxication and anti-oxidant enzymes (3). Typically, these stress responses are transiently induced in organisms exposed to toxic substances. Additionally, it is likely that they are chronically activated in the setting of degenerative disease (4), in a futile effort to minimise the ongoing macromolecular damage, although this has only been formally confirmed for progeroid syndromes (5). Thus, cytoprotective proteins are key biomarkers of organismal health, and the ability to monitor their fluctuating expression patterns promises to transform the study of drug-induced toxicity, disease pathogenesis, as well as for testing therapeutic interventions (6).

Yet, although genome-wide libraries of reporter strains have made access to gene expression patterns, including those encoding cytoprotective proteins, a routine task in model organisms, such as *E. coli, S. cerevisiae, C. elegans*, and *D. melanogaster* (7–10), these evolutionarily distant species are of limited relevance to human biology. They are particularly unsuited to addressing toxicological issues as they do not adsorb, distribute, metabolise, and excrete (ADME) compounds in the same way as humans and these and other species differences make it difficult to extrapolate results obtained with them to humans (11). Mice, are considered a much more appropriate surrogate for humans in disease and toxicology studies; not only are they more closely related, they have been genetically engineered to display ‘human-like’ ADME characteristics (12). Yet, reporter strategies have been less widely adopted by the mouse community than their counterparts working on simpler model organisms.

The marked reluctance of mouse biologists to adopt reporter mice seems to stem from concerns surrounding the fidelity of expression of reporter and target proteins. First-generation reporters, which involve random integration of reporters under the control of a promoter element, were undermined by loss of context – adjacent genetic and epigenetic elements that influence transcriptional output (13). Second-generation reporters addressed this concern by inserting the reporter element into the native gene locus in place of the target open-reading frame, and this strategy has been used to generate large collections of mouse reporter strains (14), most recently by the International Mouse Phenotyping Consortium (15). Yet this strategy still has some substantial drawbacks that limit its utility. First, protein expression is not determined solely by transcription; it is also influenced post-transcriptionally by sequences present in the target mRNA that regulate transcript stability and translatability (16–18). As the reporter cassette typically replaces some or all of these sequences, the second-generation reporter strategy remains unlikely to accurately recapitulate target protein expression patterns. Second, this strategy causes the loss of one allele and raises concerns regarding haplo-insufficiency effects. A final objection is that this strategy does not lend itself to the use of multiplexed reporters in spite of the fact that individual enzymatic, fluorescent and other reporter molecules exhibit limitations and performance could be improved by combining them (19). We recently reported (20) that the use of viral 2A technology (21) could circumvent some of these objections, and alleviate others, by allowing the expression of multiple reporters from a target gene locus while maintaining the sequences encoding the target gene.

Heme oxygenase 1 (Hmox1), a major cytoprotective enzyme found from bacteria to man, catalyses the rate-limiting step in the metabolic degradation of free heme to bile-pigments (biliverdin and bilirubin), carbon monoxide, and iron (22). It is believed that it is normally expressed at high levels only in splenic macrophages and in other tissues that degrade senescent red blood cells, including specialized reticuloendothelial cells resident in liver and bone-marrow (23). However, under conditions of hemolysis, it is dramatically induced in the parenchyma of liver and kidney to deal with the increased burden of circulating heme (24). This ensures that levels of heme, a lipophilic, freely-diffusible form of redox-active iron, are minimized, and also releases iron into the blood-stream for subsequent re-use in erythropoiesis. Mice that lack Hmox1 suffer many defects that can be explained by a failure to correctly maintain heme and iron homeostasis, including low serum iron levels and anemia associated with an accumulation of renal and hepatic iron (25); exaggerated oxidative stress; splenomegaly; and chronic inflammation. Similar symptoms were described in the only reported case to date of a patient lacking Hmox1 activity (26).

Surprisingly, Hmox1 is not induced solely in response to its natural substrate heme. Instead, it accumulates rapidly in multiple cell types, both *in vitro* and *in vivo*, in response to a bewildering array of environmental stresses and disease states (27–29). Almost without exception, *Hmox1* induction is beneficial under such circumstances. *Hmox1’*s uniquely broad responsiveness to multiple forms of tissue damage stems from the presence within its proximal promoter and distal enhancers of conserved binding sites for multiple families of stress-activated transcription factors. The most important of these factors is NFE2-related factor-2 (Nrf2) and its paralogues. However, members of the Heat Shock Family (Hsf) and NF-κB families of proteins also direct transcription of this gene (24). Its cytoprotective function is generally ascribed to its ability to convert heme to carbon monoxide and bilirubin, the first product being anti-inflammatory, and the latter an anti-oxidant (30).

Perhaps no other single cytoprotective enzyme is as important for organismal homeostasis as is Hmox1 and it is one of the most sensitive marker of organismal health available to us (24). The only *Hmox1* reporter mice described to date were first-generation reporters created by random integration of either a luciferase (31) or β-galactosidase (32) reporter genes under the control of approximately 15 kb of *Hmox1* promoter. In this paper, we describe how use of viral 2A technology has led to the development of a new and improved Hmox1 reporter mouse line. We validate the model by confirming the absence of haplo-insufficiency effects, and the fidelity of reporter expression. We provide a high-resolution blueprint for Hmox1 expression in healthy mice that suggest novel biological insights for follow-up. Finally, we exemplify how it can be used to study toxicological processes. Future work will demonstrate how the model can be applied broadly to pre-clinical research into degenerative disease.

## MATERIALS AND METHODS

### Hmox1 reporter mice

#### Construction

For the generation of Hmox1 reporter mice, a T2A-LacZ-T2A-hCG-T2A-Fluc cassette was inserted between the penultimate and STOP codons in exon 5 of *Hmox1*. The positive selection marker (puromycin resistance (PuroR)) was flanked by *FRT* sites to allow for removal after the successful generation of transgenic mice. The targeting vector was generated using BAC clones from the C57BL/6J RPCIB-731 BAC library and electroporated into TaconicArtemis C57BL/6NTac ES cell line Art B6/3.6. Positive clones were verified by PCR and Southern blot before being injected into blastocysts from superovulated BALB/c mice. Blastocysts were injected into pseudopregnant NMRI females, and the chimerism of offspring was evaluated by coat color. Highly chimeric mice were bred with C57BL/6 females mutant for the gene encoding Flp recombinase (C57BL/6-Tg(CAG-Flpe)2 Arte). Germline transmission was identified by the presence of black C57BL/6 offspring (G1). The gene encoding Flp was removed by further breeding to Flp- partners after successful verification of PuroR removal. All animal work was carried out in accordance with the Animal Scientific Procedures Act (1986) and after local ethical review. All mice were kept under standard animal house conditions, with free access to food and water, and 12h light/12h dark cycle. Data in this paper was obtained using mice of both genders and between 14 and 41 weeks of age, as specified in individual figure legends. In any given experiment, mice were age-matched to within one month of each other, unless otherwise stated.

#### Genotyping

Ear biopsies of mice 4–8 weeks old were incubated at 50 °C for 4–5 h in lysis buffer containing 75 mM NaCl, 25 mM EDTA, 1% (w/v) SDS and 100 μg/ml (39 U/mg) proteinase K (Sigma). The concentration of NaCl in the reaction was raised to 0.6 M and a chloroform extraction was performed. Two volumes of isopropyl alcohol were added to the extracted supernatant to precipitate genomic DNA (gDNA). 40 μl TE buffer (10 mM Tris, 1 mM EDTA, pH 8.0) was added to the pellet, and subsequently gDNA was dissolved overnight at 37 °C. The typical PCR sample consisted of a 25-μl volume containing 10 pmol of the primers (6395_29 5’-GCTGTATTACCTTTGGAGCAGG-3’; 6395_30 5’- CCAAAGAGGTAGCTAATTCTATCAGG-3’). Each reaction also contained 1.25 U Taq DNA polymerase (Thermo-Scientific) with 10 mM dNTPs, buffer and 25 mM MgCl_2_. The following PCR conditions were applied: 5 min, 95 °C initial denaturation; 30 s, 95 °C cyclic denaturation; 30 s, 60 °C cyclic annealing; 1 min, 72 °C cyclic elongation for a total of 35 cycles, followed by a 10-min 72 °C elongation step. All PCR protocols were developed by TaconicArtemis. PCR amplification products were analyzed by agarose gel electrophoresis.

### P21 reporter mice

These mice and accompanying genotyping protocols have previously been reported (20).

### Chemicals and γ-irradiation

All chemicals were from Sigma. Hemin was dissolved in 0.1 M NaOH, prior to adjusting pH to 7.4 with 1 M HCl. Cadmium chloride, LPS from E.coli strain O55:B5 and paraquat were all prepared in 0.9% (w/v) NaCl. Solvent was used as vehicle control in all mouse experiments. Mice were γ-irradiated in an Oris IBL 637 Cesium-137 irradiator (3 min/Gy).

### *In vivo* luciferase imaging

Imaging was performed at baseline and after chemical-dosing and/or irradiation. Reporter mice were injected *ip* with 5 μl/g body weight RediJect d-Luciferin (30 mg/ml, Caliper) and anaesthetized by isofluorane before being transferred into the IVIS Lumina II imaging chamber (Caliper) for bioluminescent imaging. Luminescent images (5 sec, f/stop 1.2, binning 4) and gray-scale images (2 sec, f/stop 16, binning 2) were acquired. Photon fluxes in Regions Of Interest (ROI) were quantified using the LivingImage (R) Software, version 4.3.1 (Caliper). ROI were defined as an oval from just below the forepaws to just above the genitals of each mice. Photon fluxes are expressed as photons/sec/cm^2^/sr. Luminescent images were rendered using the fire LUT in ImageJ, and superimposed on gray-scale photographs.

### Tissue harvesting and processing for cryo-sectioning

Mice were euthanized by exposure to rising concentrations of CO_2_. The median lobe of the liver was fixed in 10% Neutral Buffered Formalin. The proximal 2-4 cm of duodena were fixed in 4% (w/v) paraformaldehyde. All other organs, including sagittally-cut kidney, the stomach, the proximal 2 cm of the large intestine, the thymus, the spleen, the left lung lobe, the heart and the brain were fixed in Mirsky’s fixative (National Diagnostics). Tissues were stored at 4°C, formalin-fixed ones for 4 hours, Mirsky-fixed ones overnight, before being transferred into 30% (w/v) sucrose for 24 hours. Embedding was carried out in Shandon M-1 Embedding Matrix (Thermo Scientific) in a dry ice-isopentane bath. Sectioning was performed on an OFT5000 cryostat (Bright Instrument Co). With the exception of lung and brain sections, all sections were cut at 10 μm thickness with a chamber temperature of −20°C. Lung sections were cut at 12 μm thickness with a chamber temperature of −23°C. Brain sections were cut at 20 μm thickness with a chamber temperature of −23°C.

### *In situ* β-gal staining

Sections were thawed at room temperature and rehydrated in PBS supplemented with 2 mM MgCl_2_ for 5 minutes before being incubated overnight at 37°C in X-gal staining solution (PBS (pH 7.4) containing 2 mM MgCl_2_, 0.01% (w/v) sodium deoxycholate,0. 02% (v/v) Igepal CA630, 5 mM potassium ferricyanide, 5 mM potassium ferrocyanide and 1 mg/ml 5-bromo-4-chloro-3-indolyl β-D-galactopyranosidise). On the following day, slides were washed in PBS, counterstained in Nuclear Fast Red (Vector Laboratories) for 5 minutes, washed twice in distilled water and dehydrated through 70% and 95% ethanol before being incubated in Histoclear (VWR) for 3 minutes, air-dried and mounted in DPX mountant (Sigma). Images were captured using a Zeiss 12 megapixel digital camera attached to a Zeiss Axio Scope.A1 microscope, and controlled by means of the AxioVision v4.5 software (Carl Zeiss). Images were rendered in ImageJ.

### Tissue harvesting and processing for Perls Prussian Blue staining

Mice were euthanized by exposure to rising concentrations of CO_2_. Organs were fixed in Gurr buffer overnight before transfer into 70% (v/v) ethanol. The following day, organs were dehydrated and embedded in paraffin and subsequently sectioned at 5 μm thickness using a Shandon Finesse 325 microtome.

### Perls Prussian Blue staining

Sections were deparaffinised in xylene and rehydrated through a graded series of 100 – 50% (v/v) ethanol solutions, followed by immersion in distilled water. Sections were incubated for 20 min at room temperature in a freshly prepared aqueous solution containing 10% (v/v) HCl and 5% (w/v) potassium ferrocyanide. Sections were washed in distilled water, counterstained in Nuclear Fast Red for 5 min, washed in distilled water, dehydrated through 95% (v/v) ethanol and two changes of 100% (v/v) ethanol, cleared with two changes of xylene, air-dried, and mounted in DPX medium (Sigma).

### Immunoblots

Whole-cell lysates were prepared from flash-frozen organs as follows. Briefly, the organ was pulverised under liquid nitrogen with a mortar and pestle. The powder was added to Laemmli sample buffer (13 mM Tris (pH 6.8), 11% (v/v) glycerol, 0.44% (w/v) SDS,0.02% (w/v) bromophenol blue, 1.1% (v/v) 2-mercaptoethanol) supplemented with Complete EDTA-free protease inhibitors (Millipore) and HALT phosphatase inhibitors (Thermo Scientific) and vortexed vigorously. After 30 min on ice, the resulting lysate was sonicated to reduce viscosity and protein concentrations were determined using the ‘Microplate BCA Protein Assay Kit - Reducing Agent Compatible’ from Thermo Scientific, according to the manufacturer's instructions. SDS/polyacrylamide-gel electrophoresis and immunoblotting were carried out as previously described (33). Antibodies used included mouse monoclonal antibodies raised against GAPDH (clone GAPDH_71.1 (Sigma)), and β-gal (Promega), rabbit polyclonal antibodies for Hmox1 (Abcam)), goat polyclonal antibodies against firefly luciferase (Promega).

### Relative quantification of mRNA species

This was carried out by Taqman^®^ chemistry. Total RNA was isolated from cells using the RNeasy Kit (Qiagen), according to the manufacturer’s instructions. Approximately 1.0 μg of total RNA was reverse-transcribed to cDNA using the Qantitect kit (Qiagen), according to the manufacturer's instructions. The PCR mixes were prepared by mixing 1.5 μI of cDNA with 1 μI of TaqMan probe set, 10 μI of Universal PCR Master Mix (PerkinElmer Applied Biosystems) and 7.5 μI of MilliQ grade water. For the realtime PCR analysis, the following pre-designed TaqMan probe sets in solution were used: Mn00516006_m1 (Hmox1); Hs03003631 (18s ribosomal RNA) (all from PerkinElmer Applied Biosystems). A custom Taqman probe set designed against the β-galactosidase sequence used in the reporter cassette was used to measure transcript produced specifically from the reporter allele (proprietary probe and primer sequences, PerkinElmer Applied Biosystems). Data acquisition and analysis utilised the ABI PRISM^®^ 7700 sequence detection system (PerkinElmer Applied Biosystems). The relative gene expression levels in different samples were calculated using the Comparative C_T_ Method as outlined in the ABI PRISM^®^ 7700 Sequence Detection System User Bulletin #2. The expression of 18s rRNA was used as the internal control.

### Clinical chemistry

Terminal bleeds were collected in heparinized blood collection tubes (Sarstedt). Clinical chemistry assays were performed blind at the Clinical Pathology laboratory, MRC Harwell (http://www.har.mrc.ac.uk/services/pathology/clinical-chemistry).

## RESULTS

### A new model for monitoring Hmox1 expression at single-cell resolution

We created a mouse line in which an open-reading frame encoding T2A-β-gal-T2A-hCG-T2A-luciferase was inserted between the penultimate amino acid-encoding codon and the STOP codon in exon 5 of the *Hmox1* gene (Fig 1A). T2A peptides promote a phenomenon known as ribosome skipping that allows multiple proteins to be expressed from a single mRNA (21). We therefore expected that four separate polypeptides, Hmox1, β-gal, hCG, and luciferase would be expressed from this engineered allele. It is important to note that the β-gal protein was tagged with a nuclear localisation sequence to target it to the nucleus, whereas hCG is expected to be secreted as it contains a signal peptide. Finally, Hmox1 expressed from the reporter allele contains the T2A amino acid sequence at its C-terminus and is therefore distinguishable upon electrophoresis from Hmox1 expressed from the wild type (wt) allele.

**Figure 1:**
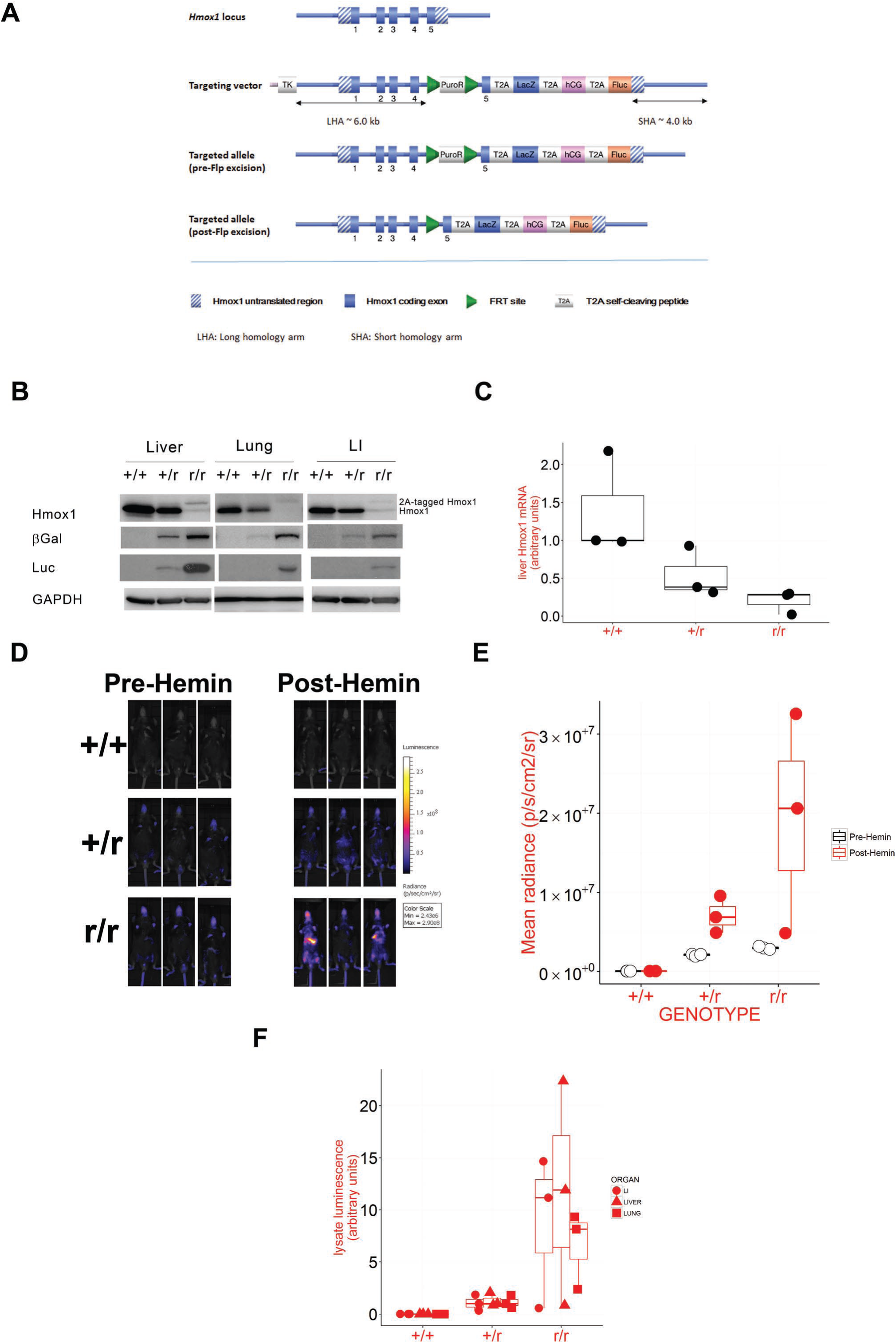
Use of T2A sequences to express β-galactosidase, luciferase, and human chorionic gonadotropin (hCG) from the endogenous *Hmox1* promoter. **A)** Schematic of the engineering strategy used to create the *Hmox1* multireporter allele. See text for further details. **B - F)** Triplicate wt (+/+), heterozygous reporter (+/r), or homozygous reporter (r/r) mice were treated with 30 mg/kg hemin and culled 18 h later. All mice were male and aged between 12 and 13 weeks except for one r/r mouse that was 7 weeks old. Note that one of the mice at the time of injection was identified as having been mis-injected. Nevertheless, data from this mouse has been included in these analyses for completeness. The data points from this particular mouse have been highlighted in the graphs of E) and F) by increased transparency. **B)** Pooled liver, lung, or large intestines (LI) whole lysates were prepared and blotted for Hmox1, β-galactosidase, and firefly luciferase. GAPDH was used as a loading control. **C)** Levels of hepatic *Hmox1* and *LacZ* mRNA relative to 18s rRNA. Data are presented as arbitrary units and have been transformed so that the median level of *Hmox1* mRNA in wt mice is set to a value of 1. The median level of *LacZ* mRNA in heterozygous reporter mice was set to 1. **D & E)** *In vivo* bioluminescence images of mice before treatment with hemin and immediately prior to sacrifice D); quantification of data E). **F)** Firefly luciferase activity per mg protein was measured in lysates prepared from lung, liver, or LI. Data are presented in arbitrary units. In order to highlight the fold-difference in activity between heterozygous and homozygous reporter mice, data have been transformed so that the median activities in organs from heterozygous reporter mice are set to a value of 1. See Fig S10A for accompanying hCG data.

We previously employed a very similar strategy to generate a p21 reporter mouse. One conclusion from this earlier work was that insertion of the reporter cassette suppressed expression of the target protein from the reporter locus, for reasons that were not further explored (20). The possibility that Hmox1 expression might similarly be compromised in the current model was a serious concern, given the severe phenotype of *Hmox1^-/-^* mice, as it would potentially disqualify it from the study of normal physiology. Therefore, to address this concern and assess the performance of the reporter allele more generally, we initially treated wild-type mice and heterozygous and homozygous reporter mice (bearing one and two *Hmox1* reporter alleles, respectively) with hemin, a classic inducer of *Hmox1*. Mice were *in vivo* imaged for luminescence pre- and post-treatment, and organs harvested for protein and mRNA analyses (Fig 1B – F).

We confirmed that four separate peptides were being correctly processed and expressed from the reporter transcript, by blotting liver, lung and large intestine lysates for Hmox1, β- gal, hCG, and luciferase proteins (Fig 1B). Specifically, we observed two bands of the predicted molecular weights for the Hmox1 expressed from the wt allele and the T2A-tagged Hmox1 expressed from the reporter allele (Fig 1B). Both β-gal and luciferase were detected as single proteins with higher expression levels in animals homozygous for the reporter allele. We could not detect hCG in any of these samples, presumably because the protein is secreted. Nevertheless, correct processing of the two polypeptides that flank it, β-gal and Luc, implies that hCG – and therefore all four polypeptides – are correctly processed.

### Heterozygous reporter mice do not display a phenotype despite suppressed expression of Hmox1

Although the 2A strategy for the expression of multiple reporters off a single allele worked well, the level of Hmox1 expression from the reporter gene was significantly lower than off its wt counterpart, as anticipated (Fig 1B). Reduced expression of Hmox1 protein from the reporter locus results, at least in part, from reduced expression of the corresponding mRNA transcript (Fig 1C). The *Hmox1* Taqman assay that we employed to investigate this issue cannot distinguish between mRNA from wt and reporter alleles, which complicates the interpretation of the data. Nevertheless, heterozygous reporter mice (one wt gene) expressed approximately half the amount of mRNA found in wild-type mice (two wt genes). Assuming that the one wt allele in heterozygous mice produces half the amount of mRNA found in wild-type mice, then these data imply that the reporter allele contributes only a small percentage of the total Hmox1-encoding mRNA found in heterozygous reporter mice. This interpretation is also consistent with the finding that the amount of *Hmox1* mRNA in homozygous reporters was much reduced in comparison with wild-type mice (Fig 1C). Consistent with compromised expression of Hmox1 protein, our homozygous reporter mice share a number of phenotypes in common with the previously described *Hmox1^-/-^* mice (25,28), including an exaggerated response to hemin. This phenomenon is evident when using a custom Taqman assay that only detects the transcript produced from the reporter locus. Using this assay, the amount of reporter transcript in hemin-treated homozygous reporter mice (which bear two reporter alleles) was considerably more than twice the amount of reporter transcript in similarly-treated heterozygous reporter mice (that bear a single reporter allele) (Fig 1C). Hyper-sensitivity to hemin is further supported by the observation that the *in vivo* (Fig 1D) and *ex vivo* (Fig 1E) luminescent signals post-hemin were far higher in homozygous than heterozygous mice. Other *Hmox1^-/-^* phenotypes observed in our homozygous reporters include splenomegaly (**Fig S1A**) and defects in iron re-utilisation (**Fig S1B**).

However, not all phenotypes are shared between the two lines with the homozygous reporter being less physiologically compromised than the complete knock-out. In particular, homozygous reporter mice were born at a reasonable, albeit lower than expected, frequency (16% observed vs 25% expected, significant at P < 0.01 by Chi-Square test) whereas the complete knockout displays almost complete perinatal lethality (25). Additionally, neither serum iron (**Fig S1C**), which is reduced in the complete knock-out (25), or serum bilirubin (**Fig S1D**) levels appear to be significantly altered in the homozygous reporter.

Crucially, heterozygous reporter mice did not display a phenotype in so far as we can judge. They display the expected response to hemin (Fig 1B). Furthermore, there was no evidence for splenomegaly or iron-loading of liver and kidneys in this line, even in older mice (**Fig S1**). We conclude that heterozygous reporter mice are suitable for studying normal physiological processes and were used in all subsequent experiments.

### Faithful expression of β-gal in the same organs, tissues, and cell types as Hmox1

An essential attribute of any reporter system is that the pattern of expression of the reporter proteins faithfully mimics that of the target. We anticipated that this would be the case in our model, as all the coding and regulatory elements that might control Hmox1 expression are retained in the reporter allele. Unfortunately, the true expression pattern for Hmox1 in healthy adult mice has not been systematically characterized, as evidenced by the lack of data curated in Mouse Genome Informatics (MGI; http://www.informatics.jax.org). Although the absence of such data highlights why reporter models such as ours are necessary, it does present problems when attempting to validate them. We therefore initially documented β-galactosidase expression patterns in spleen, liver, and kidney, as detailed expression data is available for Hmox1 in these three organs under a variety of conditions.

For example, Hmox1 activity in healthy animals is believed to reside in splenic and hepatic macrophage populations (22–24), and in proximal tubular epithelial cells of the kidney (34,35). Consistent with this, we consistently detected populations of β-galactosidase-positive cells in spleen and liver sections (Fig 2A). Based on size and morphology, positive cells seemed to belong to the macrophage population. We also found renal expression of Hmox1 in tubules – but not glomeruli – of the cortex (Fig 2A).

**Figure 2:**
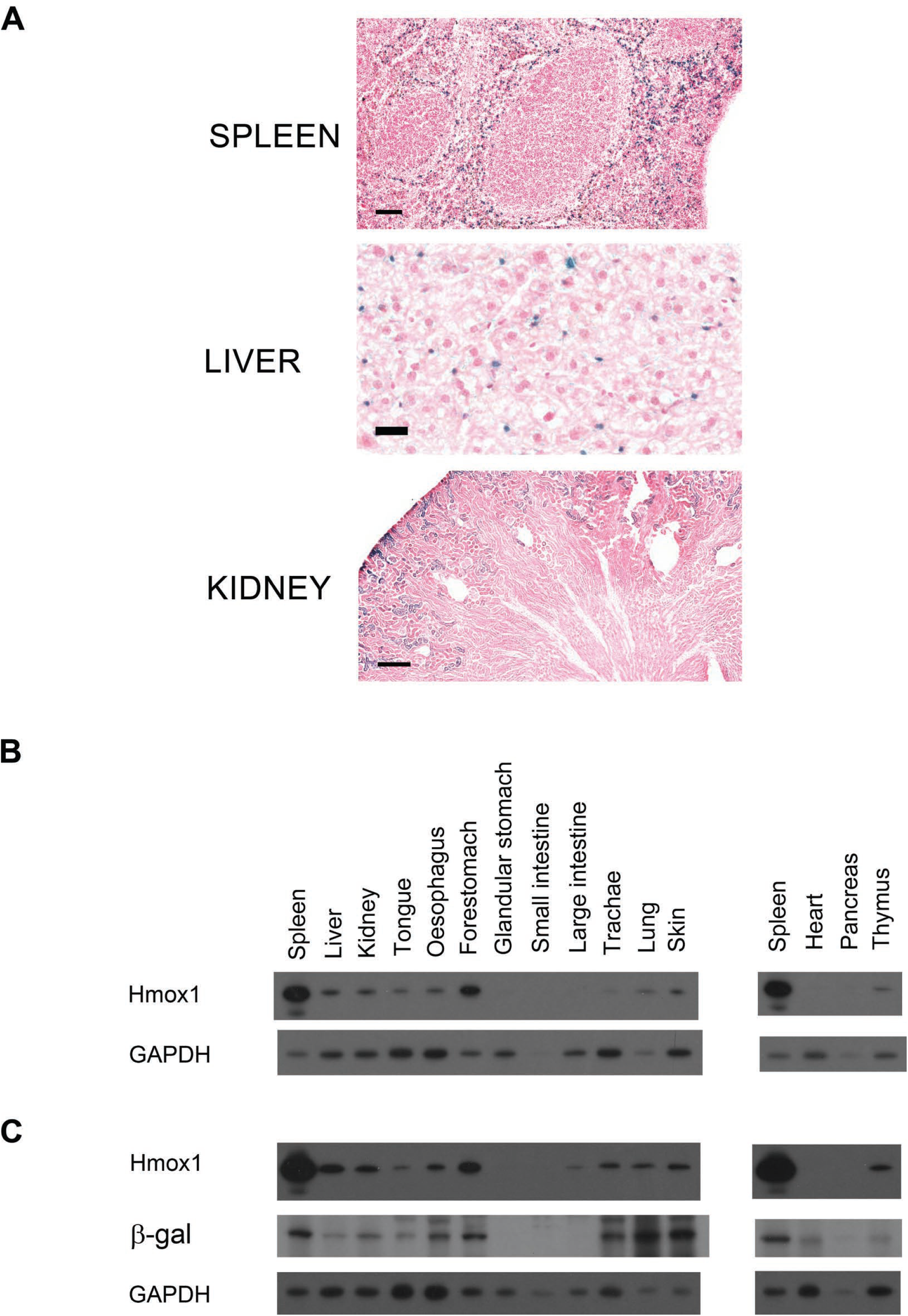
*In situ* β-galactosidase activity in spleen, liver, and kidney of control reporter mice. **A)** Representative spleen, liver, and kidney sections from male mice aged between 10 and 14 weeks of age. Scale-bar = 100 μm (spleen), 50 μm (liver), 250 μm (kidney). See Fig S2 for data on the gastro-intestinal tract; Figure S3 for trachea, lung and skin data; Figure S4 for thymus, pancreas, and heart data. **B & C)** Pooled organ lysates (n = 4) from untreated mice, wt (B) or reporter (C), were probed for Hmox1 and β-galactosidase. GAPDH was used as loading controls. The β-galactosidase blot was negative in wt mice and is not shown.

Exposure of mice to hemin induces *Hmox1* in hepatocytes and renal tubules, whilst leaving levels unaltered in the spleen (23,35). In excellent agreement with these data, hemin induced reporter activity in hepatocytes and cells of the renal tubule; moreover, the staining pattern was unchanged in the spleen after treatment (Fig 3B).

**Figure 3:**
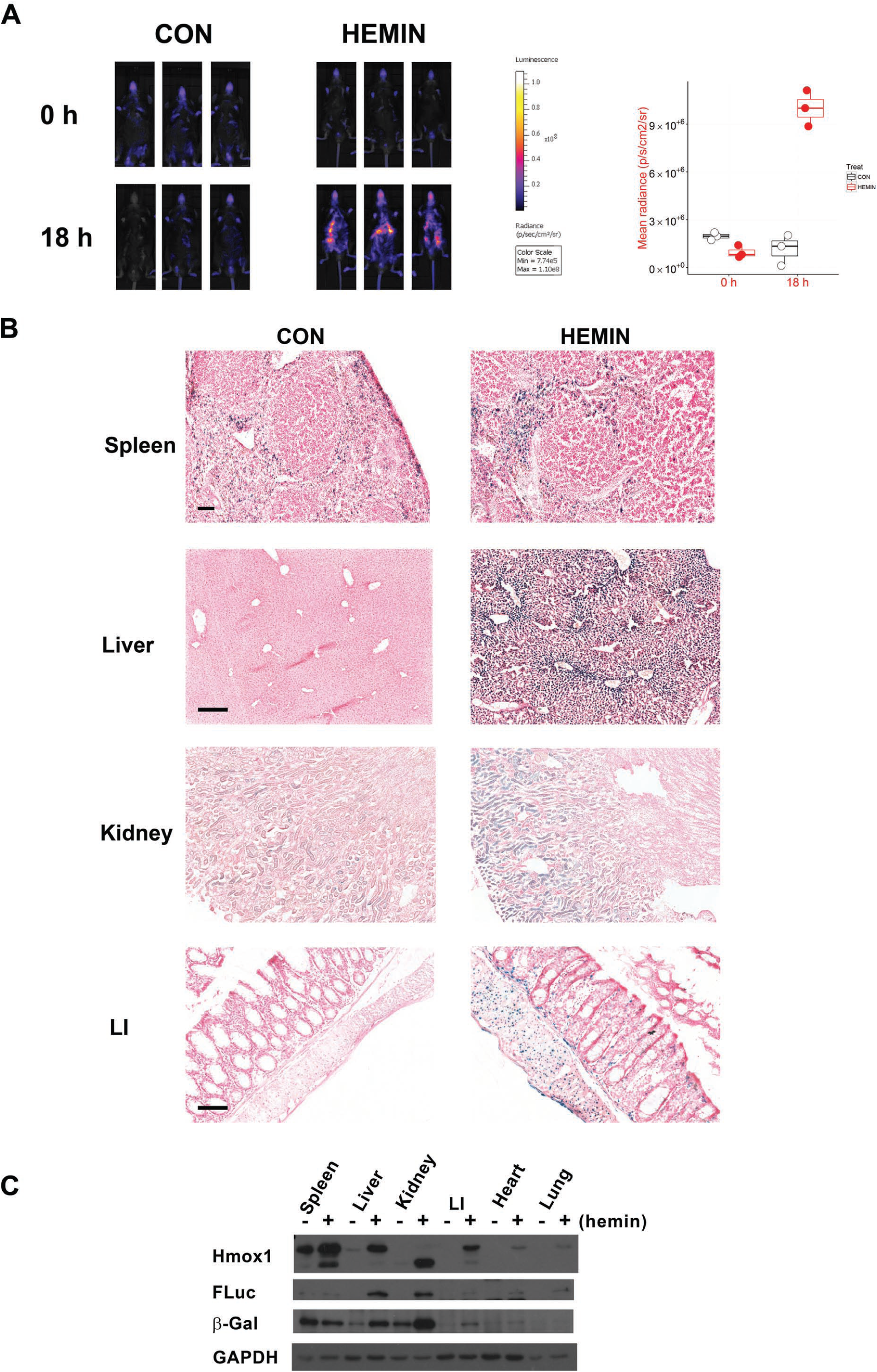
Hemin induces Hmox1 reporters in liver and kidney cells. Triplicate reporter mice were treated with 30 mg/kg hemin and culled 18 h later. All mice were male and aged between 19 and 20 weeks except for one hemin-treated mouse that was 14 weeks old. **A)** Mice were imaged at baseline and, again, just before necropsy. The accompanying graph quantifies the luminescent signal. **B)** Spleen, liver, kidney, and large intestine (LI) sections from control and hemin-treated mice were *in situ* β-galactosidase stained. Representative images are shown. Scale-bar = 250 μm except LI (100 μm). **C)** Pooled organ lysates (*n* = 3) were probed for the indicated proteins. GAPDH was used as a loading control. See Figure S10B for accompanying hCG data.

Finally, CdCl_2_ is a nephro- and hepatoxin that potently induces Hmox1 activity in both organs (36–38). We observed massive induction of reporter activity in renal tubules and liver hepatocytes, in agreement with these previous reports (Fig 4B). Strikingly, this experiment included mice of both genders and the data hint at sexual dimorphism in expression of Hmox1 in control and treated mice (Fig 4).

**Figure 4.**
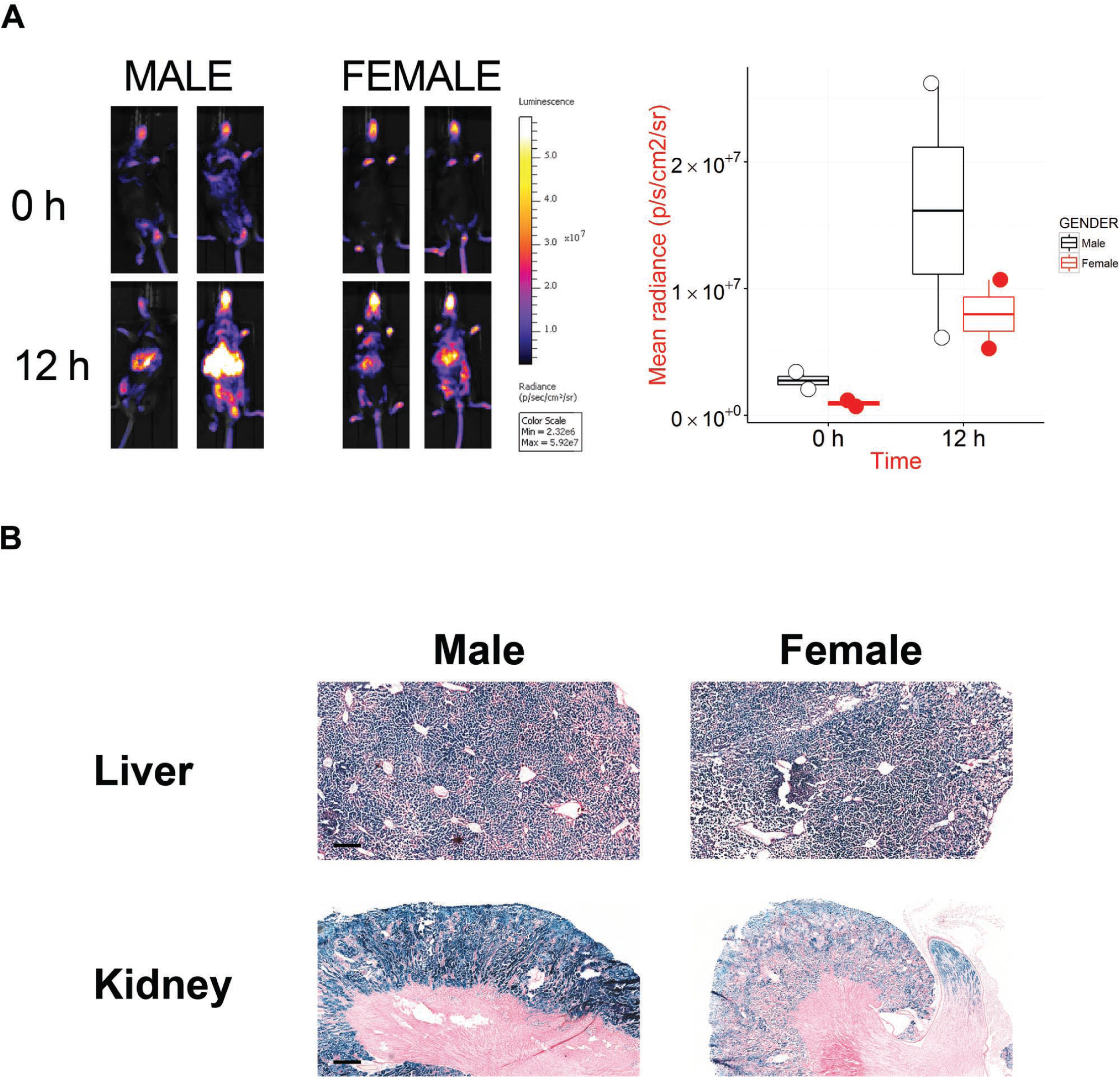
Cadmium chloride causes sexually dimorphic induction of Hmox1 reporter signals in liver and kidney. Two male and two female mice were treated with 4 mg/kg cadmium chloride (Cd). **A)** Bioluminescent images were obtained both before and 12 h after exposure to Cd. Luminescence is quantified in the accompanying graph. **B)** Liver and kidney sections were stained for β-galactosidase activity. Representative images are shown. Scale-bar = 250 μm (liver) and 500 μm (kidney).

Taken in total, these data validate our reporter system as a *bona fide* marker for Hmox1 protein expression, insofar as the limited pre-existing independent data allow us to make this assessment.

### Novel Hmox1 expression patterns identified through high-resolution reporter analysis

To extend our currently limited understanding of Hmox1 expression patterns in healthy mice beyond spleen, liver and kidney, we performed *in situ* β-galactosidase staining on a comprehensive selection of additional mouse organs (**Figs S2 – S4**). Unexpectedly, we noticed that reporter protein was expressed at high levels in those barrier tissues at the interface between the internal and external environment that have evolved to provide mechanical and chemical protection to the organism.

In the alimentary tract for example, reporter expression is expressed solely in the stratified, cornified, epithelial layer that extends from the tongue, through the oesophagus, and down to the juncture between the fore- and glandular stomach (**Fig S2**). At this juncture, the structure and function of the epithelial lining changes from stratified and protective to simple columnar specialized for adsorption and reporter positivity ends (**Fig S2**). Similarly, in the respiratory system, the protective lining of the trachea and bronchi are highly and modestly positive, respectively (**Fig S3**). Finally, the stratified, cornified epithelial layer covering the skin, a major point of defense against the external world, was highly positive (**Fig S3**).

By way of contrast with spleen, liver, and kidney, other internal organs, such as heart, pancreas, and thymus (**Fig S4**), did not express detectable levels of β-galactosidase.

Given the unexpected nature of these findings, we tested the prediction that Hmox1 would be found in multiple barrier tissues by blotting lysates prepared from pertinent organs of wt mice (Fig 2B). Although this approach only provides us with a crude, average measure of Hmox1 across all cell types present in an organ, there are no authenticated IHC-ready antibodies that could be used to provide higher-resolution date. The results were in excellent agreement with predictions. For example, we confirmed that the respiratory system and skin expressed significant amounts of Hmox1 (Fig 2B). It was particularly gratifying to note that the expression pattern of Hmox1 throughout the alimentary tract confirmed in all important respects to our predictions; tongue, oesophagus, and forestomach were positive for Hmox1 whereas small- and large intestines were essentially negative (Fig 2B).

Collectively, these data highlight the power of our reporter technology to chart target protein expression. They also represent the most comprehensive characterization to data of Hmox1 expression in any adult mammal and suggest novel biological insight into its physiological role *in vivo* (see Discussion).

### Correlation of target Hmox1 protein levels and reporter levels

Because reporter proteins and target proteins will generally have different half-lives, and their relative half-lives may vary from organ to organ, it was not clear how well-correlated levels of luciferase, β- galactosidase, and Hmox1 proteins would be. In fact, with one notable exception, levels were well-correlated across an extensive panel of organs obtained from untreated reporter mice (Fig 2C). The major discrepancy was that levels of both reporter proteins were lower in spleen extracts than would be predicted based on the overall trend. We also examined the correlation between target and reporters in mice treated with hemin (Fig 3C), which again revealed a high degree of correlation. There is some indication, however, that the dynamic range of β-galactosidase is compressed compared to Hmox1, perhaps reflecting its long halflife.

### Inflammation and oxidative stress induce reporter expression

The major pathophysiological states leading to increased expression of Hmox1 are inflammation and oxidative stress (29,30). We modelled inflammation by exposing our reporter mice to LPS (Fig 5). The best-described effect of inflammation on Hmox1 expression is of a significant but transient induction of Hmox1 in hepatocytes as part of the acute phase response (39), although the increase in protein is much more modest (40,41). Consistent with this, we observed at best a very modest increase in the *in situ* β-galactosidase signal in liver sections of treated reporter mice. The modest fold of induction was confirmed by blotting lysates from these same control and LPS-treated mouse livers (**Fig S5**).

**Figure 5:**
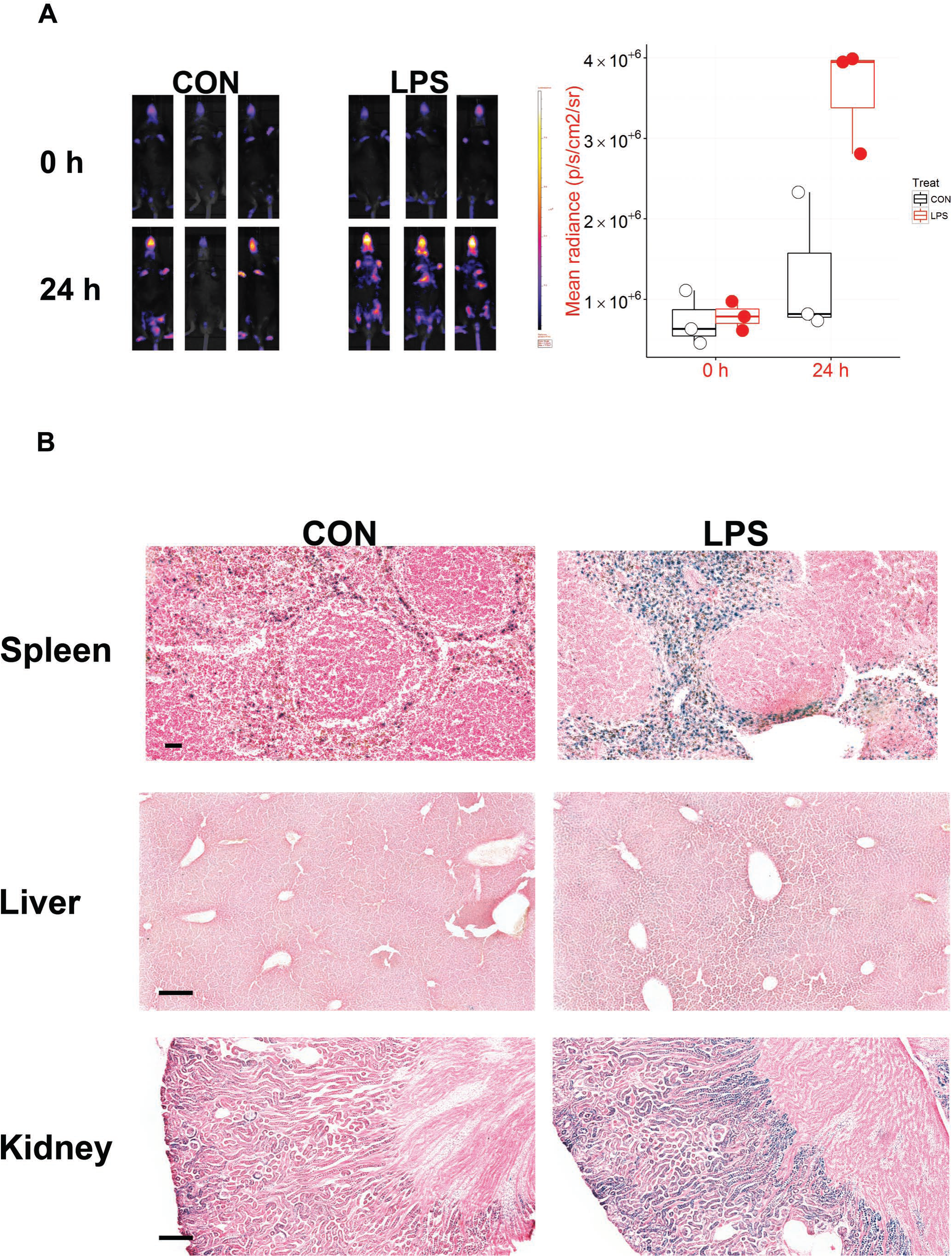
LPS induces reporter activity in spleen and kidney, but not liver. Triplicate reporter mice were treated with vehicle (PBS) or 1 mg/kg LPS. Two male and one female mice were used per treatment group. All mice were aged 18 weeks. **A)** Mice were imaged for bioluminescence at baseline and just before necropsy (24 h post-treatment). **B)** All livers and kidneys were stained for β-galactosidase activity. Representative images are shown. Scalebar = 250 μm. The corresponding sections from tongue, oesophagus, LI, heart, and lung can be found in Figure S6. Liver lysates were prepared and blotted for Hmox1, β-galactosidase and can be found in Figure S5.

Although LPS had a modest effect of hepatic expression of Hmox1 and reporter expression, it caused a significant elevation in β-galactosidase expression in both splenic macrophages and tubular epithelial cells of the kidney cortex (Fig 5). A striking finding that has not been reported previously is that muscle cells in multiple anatomical locations, including the tongue, oesophagus, large intestine, responded to LPS by inducing Hmox1 protein, as inferred from *in situ* β-galactosidase staining (**Fig S6**). Cardiomyocytes responded in a similar manner (**Fig S6**). Muscle cells seem to be a major source of Hmox1 protein as hemin, to our surprise, induced β-galactosidase activity in the smooth muscle cell layers surrounding the large intestine (Fig 3B).

To model oxidative stress, reporter mice were injected with paraquat, a redox cycling compound that produces superoxide and hydrogen peroxide in large amounts (42). It is exceeding toxic to the pulmonary system where it is concentrated by the polyamine transporter system in epithelial cells lining the lung, although it also causes serious damage to the kidney, liver, and heart (42–44). Consistent with these pathological findings, there was a massive increase in β-galactosidase staining in lungs from treated reporter mice and this was restricted to alveolar epithelial cells (**Fig 6B**).

These data are important because they reveal that our reporter mice can be used to study the numerous disease states associated with inflammation and oxidative stress.

**Figure 6:**
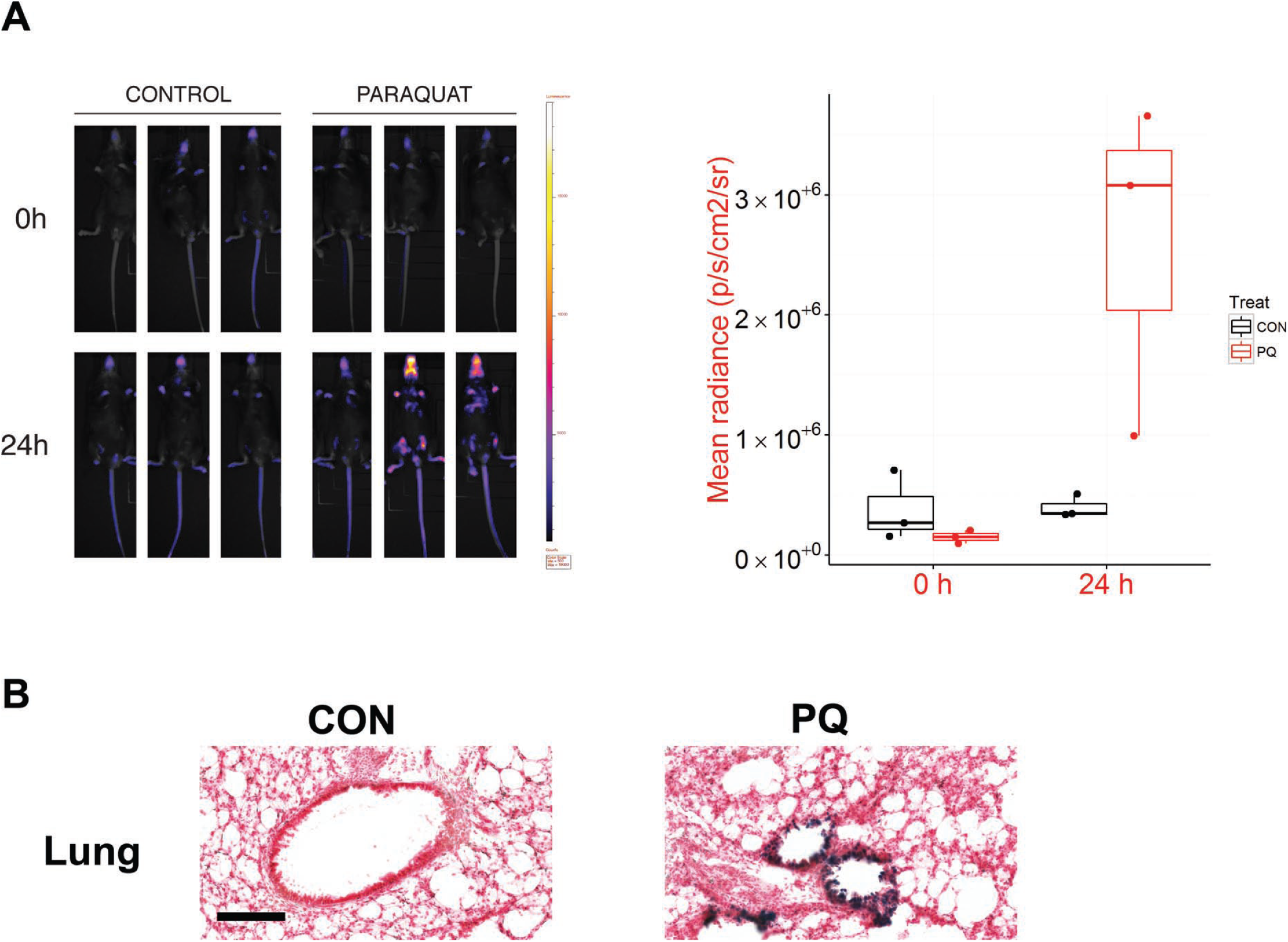
Paraquat induces reporter activity in alveolar epithelial cells. Triplicate male reporter mice aged between 12 and 14 weeks of age were treated with vehicle (PBS) or paraquat (30 mg/kg). **A)** Mice were imaged for bioluminescence at baseline and just before necropsy (24 h post-treatment). **B)** All lungs were stained for β-galactosidase activity. Representative images are shown. Scale-bar = 50 μm.

### Monitoring of luminescence from internal organs in living mice

We confirmed that the luciferase reporter facilitated the non-invasive luminescent imaging of *Hmox1* induction. For example, exposure of reporter mice to hemin (Fig 3A), cadmium (Fig 4A), LPS (Fig 5A), and paraquat (Fig 6A) resulted in significantly increased bioluminescence over that observed in control counterparts. Unlike our *p21* reporter model (20), bioluminescence from visceral organs is visible in the Hmox1 model as the signal is not over-whelmed by a signal emanating from the skin. However, resolution of luminescent signals is still low and although it is possible to make an informed estimate as to the possible identities of the organ(s) involved, unambiguous identification is still difficult.

### Ionizing radiation does not induce acute oxidative stress or inflammation

Most of the agents and states known to induce Hmox1 protein damage lipids and proteins rather than DNA. Thus, the *Hmox1* reporter line should provide orthologous and complementary data to that provided by our *p21* DNA damage reporter. To confirm that DNA damage does not acutely activate the Hmox1 system *in vivo*, we exposed Hmox1 reporter mice to 4 Gy of IR.

This did not result in elevated *in vivo* bioluminescence measurements (Fig 7A). More convincingly, no increase in *in situ* β-galactosidase signal, above that of control mice, was observed *post-mortem* (Fig 7B & **Fig S7**). By way of contrast, a massive increase in the β- galactosidase signal was observed in similarly-treated p21 reporter mice (Fig S8). In fact, in the IR-sensitive large intestine, the signal was already saturated even at the lowest dose of IR tested (1 Gy).

**Figure 7:**
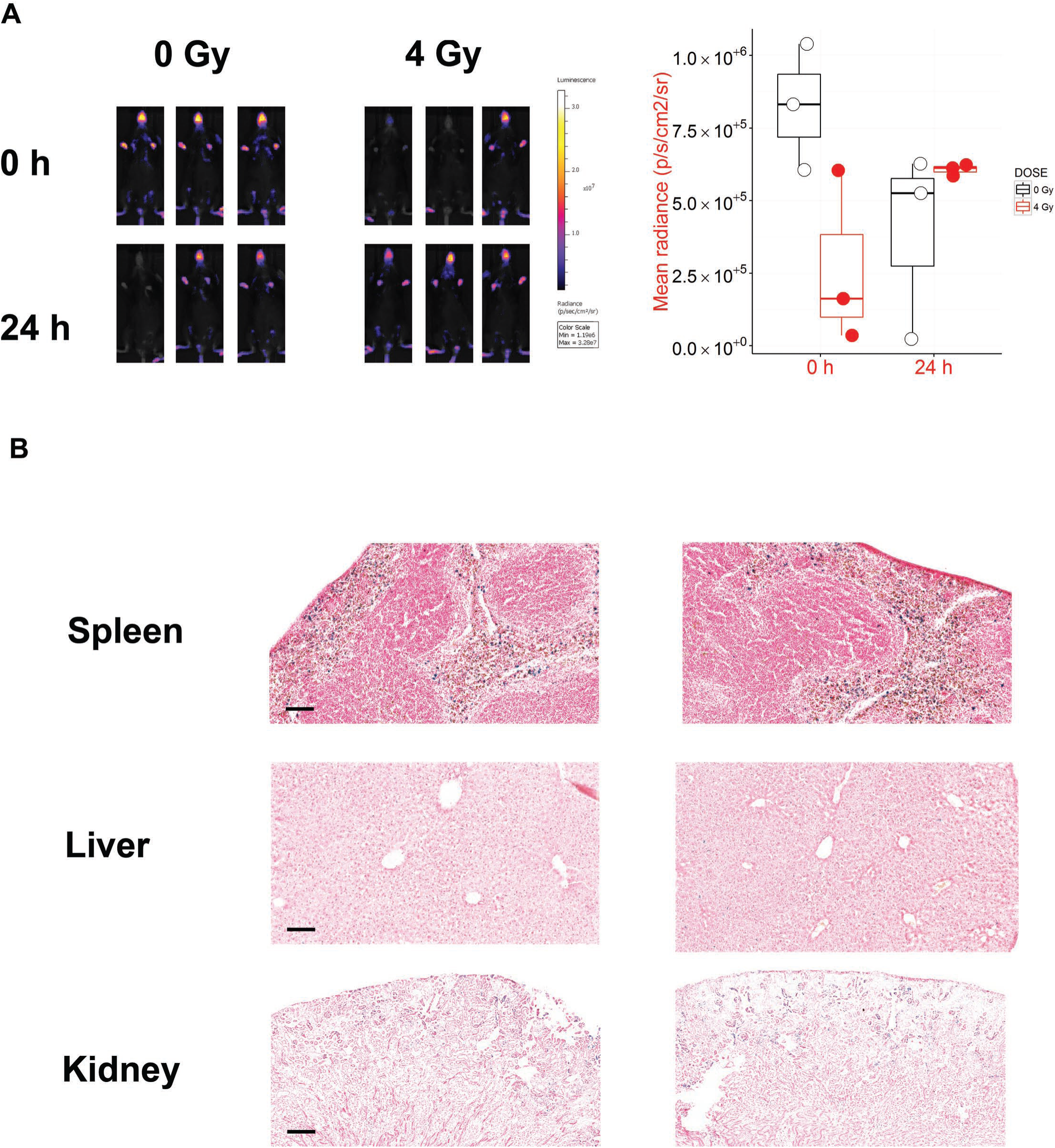
Ionizing radiation (IR) does not alter bioluminescence in reporter mice. Triplicate reporter mice were sham-irradiated (0 Gy) or exposed to 4 Gy of IR. All mice were female and aged between 17 and 21 weeks of age. **A)** Mice were imaged for bioluminescence at baseline and just before necropsy (24 h post-irradiation). The accompanying graph quantifies the radiance from each mouse. **B)** Spleen, liver, and kidney sections were stained for ß- galactosidase activity. Scale bar = 100 μm (spleen and liver) and 250 μm (kidney). See Figure S7 for accompanying *in situ* β-galactosidase staining of LI, lung, and heart.

### Acetaminophen (APAP) toxicity predicted by the Hmox1 reporter

Finally, to exemplify how this model can be used in a toxicological sense, we examined whether our *Hmox1* reporter would correctly identify *N*-acetyl cysteine as an antidote to APAP poisoning. APAP, exposure to which induces Hmox1 activity (45), is a potent hepatotoxin ((46,47) and Fig 8A) as a consequence of its metabolic conversion to a highly thiol-active electrophile in hepatocytes; the thiol-containing prodrug *N*-acetyl cysteine is an effective antidote ((48,49) and Fig 8A). Gratifyingly, APAP not only induced reporter activity in hepatocytes, but it did so specifically in the centrilobular zone (Fig 8B), as expected (46). As we have previously reported, the cells identified as Hmox1-positive surrounded a core of necrotic cells (45). Importantly, co-administration of *N*-acetyl cysteine reduced the reporter signal, and prevented necrosis of centrilobular hepatocytes (Fig 8B). These now-surviving cells, however, displayed a mild induction of Hmox1, as assessed by β-galactosidase activity (Fig 8B), indicating that they may still be subject to some degree of damage.

**Figure 8:**
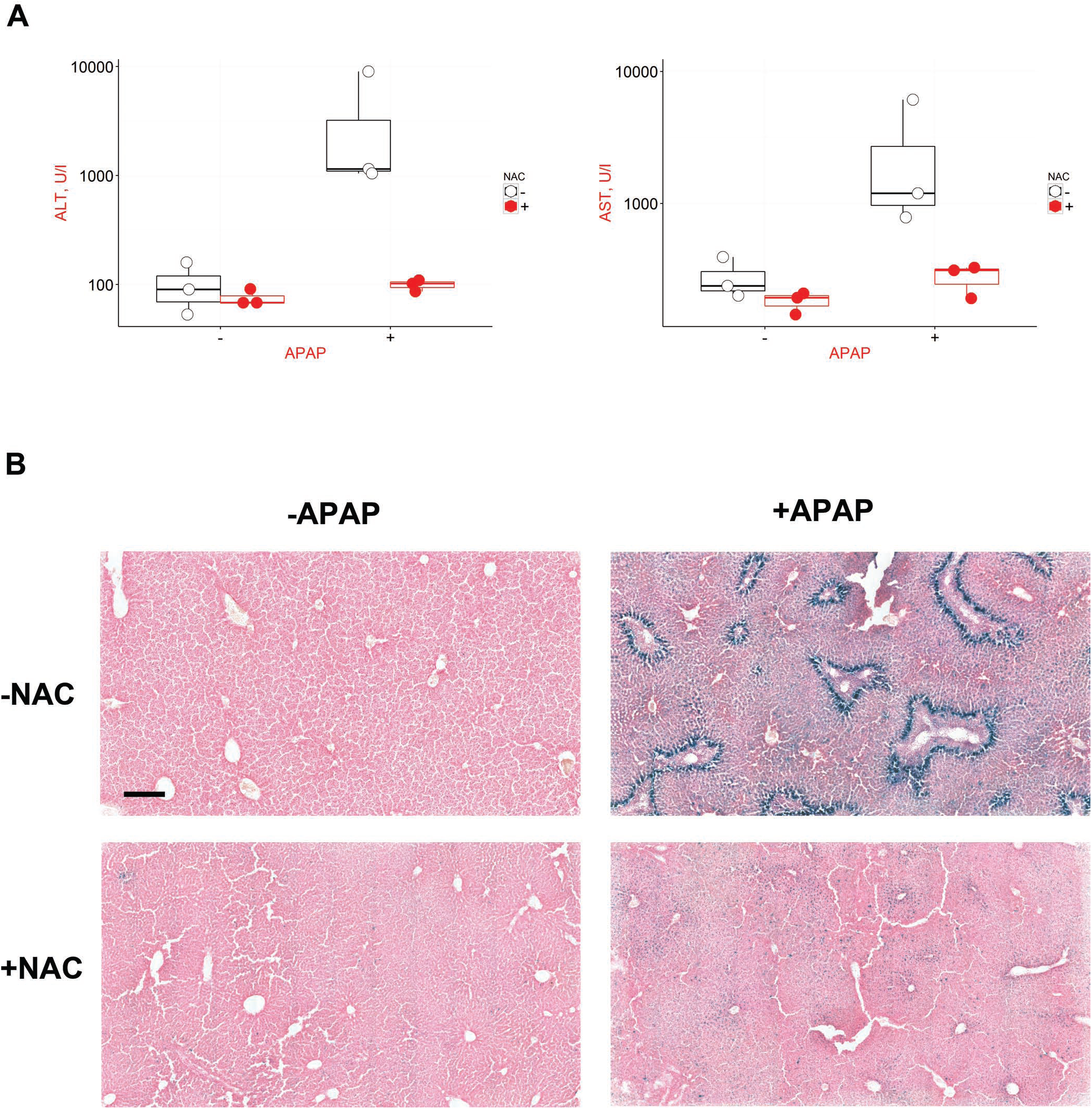
APAP induces reporter activity in the liver that is mitigated by the antidote NAC. Triplicate reporter mice were treated with vehicle, 300 mg/kg APAP, 300 mg/kg NAC, or a combination of 300 mg/kg APAP + 300 mg/kg NAC, and necropsied 24 h later. Each treatment group contained two male and one female mouse and mice were between 9 and 10 weeks of age. **A)** ALT and AST activities in plasma obtained at necropsy. **B)** All livers were sectioned and stained for β-galactosidase activity. Representative images are shown. Scale-bar = 250 μm. In addition, mice were imaged for bioluminescence before treatment and just prior to necropsy – data can be found in Figure S9.

The accompanying *in vivo* bioluminescence data did not display a convincing difference in signal between treatment groups (**Fig S9**). This is presumably due to the fact that with APAP-elicited β-galactosidase-positivity is restricted to a small fraction of hepatocytes.

### hCG measurements are unreliable

We included a hCG reporter in our reporter cassette in the hope that it would allow us to infer Hmox1 expression in serial blood samples. However, as illustrated by the data in **Fig S10**, serum levels of hCG measured with a commercial ELISA only rarely exceeded the limit of detection of the assay and were unreliable as a result. Therefore, this reporter was not pursued any further.

## DISCUSSION

In this article, we describe a method for assessing Hmox1 expression across all cell lineages in mice. It was developed to allow expression to be monitored non-invasively over extended periods of time (bioluminescent imaging), whilst permitting single-cell resolution analyses *post-mortem* (β-galactosidase staining). The goal of this first study was to validate and rigorously assess the performance of the model, provide a blue-print for Hmox1 expression in healthy mice, and exemplify its use in toxicology.

### Model performance – reduced expression of Hmox1 protein from the reporter locus

By using viral 2A sequences, we hoped that our reporter strategy would maintain expression of Hmox1 protein in order to reduce the possibility of undesirable haplo-insufficiency effects resulting in disturbed physiology. In the event, expression of Hmox1 protein was suppressed from the reporter allele. Suppression of Hmox1 protein occurs at least in part because, for reasons that are unclear, steady-state levels of reporter transcript were considerably lower than expected based on the level of transcript from the endogenous allele. In spite of this suppression, heterozygous reporter mice did not display any overt phenotypes and therefore seem suitable for investigating normal physiology. This conclusion is supported by the finding that many healthy humans only possess one functional Hmox1 allele (ExAC consortium: http://exac.broadinstitute.org/gene/ENSG00000100292).

### Model performance – fidelity and accuracy

Contrary to existing systems, our reporters are expressed from the endogenous locus and as part of the wild-type mRNA. As gene expression is not only controlled transcriptionally by DNA elements both proximal and distal, such as promoters and enhancers (50), but also post-transcriptionally by sequences resident in the target transcript (51–55), this is a major advantage of our reporter strategy over the current state-of-the-art. Our system should respond not only to transcriptional regulation but should also reflect important post-transcriptional regulatory mechanisms, including control of mRNA stability by miRNA and RNA-binding proteins and control of rate of translation (16–18). In fact, the only absolute limitation on our strategy is that it cannot report on post-translational control of gene expression, as the reporters are not fused to the gene product.

We validated the fidelity of reporter expression in a number of ways. Firstly, although Hmox1 expression patterns *in vivo* are not well-defined, we generated benchmark Hmox1 expression data across a panel of organs in control and hemin-treated mice and confirmed that reporter and target (Hmox1) protein levels were well correlated in general. Secondly, these population-level analyses were extended in the case of spleen, liver, and kidney by confirming at the single-cell level that reporter protein was expressed in the same cell types as the target. However, in general, high-resolution Hmox1 expression data is lacking and concerted efforts will need to be made to formally validate the new expression patterns described herein (see below). In the absence of antibodies validated for immunohistochemistry, this issue might best be addressed using newer tools, such as imaging mass spectrometry (56).

As with any reporter strategy, the stability of the reporters and target proteins can vary; thus reporter expression is to some extent a 'distorted' measure of target protein abundance. This presumably explains the lack of perfect correlation amongst expression levels for reporters and Hmox1 across organs and conditions.

### Identification of new expression patterns

Gene expression patterns can provide novel biological insights for subsequent follow-up. We made two particularly striking observations about the expression of Hmox1 that hint at new biology. First, healthy mice constitutively express Hmox1 in barrier tissues lining the alimentary, respiratory, and integumentary systems. Much of the positive barrier tissue is comprised of *Stratum corneum (Sc)*, which is traditionally considered to be ‘dead’ and to be incapable of engaging in biochemistry. The perception that such tissue is non-viable might cast doubt on the significance (or even veracity) of our inference that Hmox1 is expressed in such a location. However, note that Nrf2, the major factor governing *Hmox1* transcription, is already known to drive expression of Sc-restricted genes, such as *Slpi* and *Sprr2d* (57,58), and is therefore well placed to express *Hmox1* in such a spatially-restricted manner. Moreover, biochemistry is possible in the Sc; for example citrullination of fillagrin (59).

What role might Hmox1 play in the Sc? The *Sc* is comprised of corneocytes embedded in lipids that are susceptible to peroxidation arising from exposure to UV irradiation (60). Bilirubin, the end-result of Hmox1 enzymatic activity, is a lipid-soluble antioxidant and might be expected to exercise a protective role by neutralizing peroxidation. Indeed, it has been reported to be produced in the skin specifically in the *Sc* (61). On this basis, we hypothesize that Hmox1 might be responsible for *in situ* production of this chemical at barrier tissues. This leads to the testable prediction that Hmox1 deficiency might contribute to skin and other diseases as a consequence of barrier dysfunction. Significant biological effort will be required to follow up on this initial observation.

A second surprising finding was that muscle cells seem to be a major source of Hmox1 in stressed animals. Hemin induced Hmox1 protein in smooth muscle cells lining the large intestine, for example. Similarly, inflammation induced Hmox1 in muscle tissue of several organs. We suggest that this is because myoglobin, the second most abundant hemeprotein after haemoglobin, is restricted to muscle cells. It makes sense to express Hmox1 in cell types containing large amounts of heme, especially when the resulting cytoprotective products are highly mobile and can diffuse to adjacent tissues.

### Application to study human degenerative diseases

In this work, we used the example of APAP poisoning to exemplify the toxicological applications of our model. With this tool, in combination with our similar p21 reporter line, it should be possible to identify potential toxins amongst drugs, industrial- and other chemicals of interest to human health and decipher their mode-of-action.

However, we anticipate that a major future use of this model and our previously described *p21* model will be pre-clinical research into degenerative conditions. Many such conditions, including Alzheimer’s and Parkinson’s disease are associated with DNA damage or oxidative stress (4) and it is reasonable to speculate that this will lead to exacerbated stress responses. However, with few exceptions like progeroid syndromes (5), this hypothesis has not been rigorously examined due to a lack of appropriate stress-monitoring tools. Our models provide a solution to this problem and will enable researchers to monitor stress responses *in vivo* as the disease progresses. We note that there is an unmet need for prognostic biomarkers of degeneration that can improve the ability of pre-clinical research to correctly predict future clinical benefit (62). Were reporter signals to be exacerbated to a significant extent in mouse models of degenerative disease, then this would be a very attractive approach to this problem, especially so if they become elevated at or before the prodromal stage.

### Future development of technology

We believe that our 2A reporter strategy is a significant improvement on the current state-of-the-art, represented by the approaches taken by the Velocigene project (14) and the more recent International Mouse Phenotyping Consortium (15). If it is to reach its full potential, however, there are a number of improvements that need to be made. Firstly, although data from the ExAC consortium predicts that 80% of genes will not be found to suffer from haplo-insufficiency (63), it remains desirable to avoid the multireporter DNA cassette suppressing expression of the target protein. So far, the initial cassette described here, in our p21 model (20), and also used in an IL-6 reporter model (unpublished data) has this suppressive effect, which therefore appears to be an intrinsic feature. It should prove possible, however, to optimise the type and number of reporter genes incorporated into the cassette so as to avoid target protein suppression. Intuitively, shorter reporter cassettes should be less disruptive of gene expression than longer ones. Preliminary results indicate that the shorter GFP reporter is less disruptive than is the larger β-galactosidase (Ding, McMahon, Henderson, and Wolf – unpublished data) and perhaps should be used in future. A further advantage of this approach is that it would allow one to produce mice reporting on multiple genes by using fluorescent reporters with distinct spectral characteristics. One caveat to this is that concerns have been expressed over the comparatively lower sensitivity of GFP compared to β-galactosidase (13). Additionally, the hCG reporter could be removed as in our hands it has not proven useful.

A second improvement relates to the natural stability of β-galactosidase that compresses the dynamic range of the reporter and reduces its sensitivity; it would be beneficial to destabilize it or any other reporter.

Finally, a third improvement relates to the choice of non-invasive reporter. *In vivo* bioluminescent imaging of luciferase is still plagued by technical difficulties and concerns, including, but not limited to, low resolution, as discussed elsewhere (64–66). It may be better to use other imaging modalities, such as positron emission tomography or magnetic resonance imaging that are capable of non-invasive imaging of deep tissues at higher resolution.

